# A stress-activated neuronal ensemble in the supramammillary nucleus produces anxiety-like behavior in male mice

**DOI:** 10.1101/2025.01.15.633288

**Authors:** Jinming Zhang, Kexin Yu, Junmin Zhang, Yuan Chang, Xiao Sun, Zhaoqiang Qian, Zongpeng Sun, Yanning Qiao, Zhiqiang Liu, Wei Ren, Jing Han

## Abstract

Anxiety is a prevalent negative emotional state induced by stress; however, the neural mechanism underlying anxiety is still largely unknown. We used acute and chronic stress to induce anxiety and test anxiety-like behavior; immunostaining, multichannel extracellular electrophysiological recording and Ca^2+^ imaging to evaluate neuronal activity; and virus-based neuronal tracing to label circuits and manipulate circuitry activity. Here, we identified a hypothalamic region, the supramammillary nucleus (SuM), plays important role in anxiety-like behavior. We then characterized a small ensemble of stress-activated neurons (SANs) that are recruited by stress. These SANs respond specifically to stress, and their activation robustly increases anxiety-like behavior in male mice. We also found that ventral subiculum (vSub)-SuM projection but not dorsal subiculum (dSub)-SuM projections encode anxiety-like behavior and that inhibition of these vSub-SuM projections has an antianxiety effect. These results indicate that the reactivation of stress-activated supramammillary cells and relevant neural circuits are important neural processes underlying anxiety-like behavior.

## Introduction

Anxiety is a fundamental negative emotion observed in almost all mammal species. Long-lasting and uncontrollable anxiety often leads to several mental disorders, anxiety disorders, and even depression[5]. Recent studies have shown that the supramammillary nucleus (SuM), a part of the hypothalamus, regulates sleep[6], memory[7, 8], novelty exploration[7, 9], social memory[10, 11], neurogenesis[12], consciousness[13], locomotor activity[14], and theta oscillations in the hippocampus[15]. Projections from the SuM to the hippocampus have been largely studied and were found to modulate either episodic memory[7] or social memory[10, 11] in depending on the subregion of the hippocampus targeted. Although the SuM is located near the mammillary nucleus, a key region implicated in emotion regulation via the Papez circuit, its role in regulating emotion has been explored only superficially, without in-depth investigation. Despite some discussions of this role of the SuM, no consistent conclusion has been reached thus far[12, 16, 17].

Activity-dependent activation of cells have been studied in many brain areas[18, 19]. Tagged cells react to specific stimuli, such as conditional stimuli[20, 21], pain[22], food[23], and even peripheral inflammation[24], and mediate the storage and retrieval of relevant memories. The manipulation of those believed memory-associated cells can alleviate neurodegenerative diseases[25] or inflammation[24]. Naturally, this has led us to consider whether there is special neuronal ensemble that play roles in regulating anxiety or anxiety-like behaviors.Click or tap here to enter text. Recent studies have focused on the role of the hippocampus and related neuronal afferents and efferents[27–29]. The dorsal part of the hippocampus mainly contributes to cognition, whereas the ventral hippocampus is often associated with emotion[30, 31]. Although the ventral hippocampus□hypothalamus circuit was reported to modulate anxiety[27, 28], it is still unknown whether the SuM is part of this regulatory circuit. Although the SuM sends and receives dense neuronal projections, few studies have focused on its afferents or its ability to modulate behavior and emotion[32].

In this study, we hypothesize that stress can recruit a special neuronal ensemble that exclusively encodes anxiety. To test this hypothesis, we first used multiple methods to assess whether the SuM responds to acute or chronic stress. The activity of the SuM was chemogenetically manipulated, and anxiety-like behavior in rodents was tested. We subsequently employed the targeted recombination in active populations (TRAP) strategy to label and manipulate stress-activated neurons in the SuM in terms of anxiety-like behavior. After demonstrating how the SuM modulates anxiety, we sought to identify upstream brain areas that may contribute to supramammillary function. We also examined the functions of neuronal projections from the ventral subiculum (vSub) to the SuM using fiber photometry to measure calcium dynamics, as well as chemogenetic manipulation. These results allowed us to characterize a previously unreported role of SuM in regulating anxiety-like behavior. In addition, we showed that projections from the vSub but not the dorsal subiculum (dSub) to the SuM govern chronic stress-induced anxiety-like behaviors.

## Results

### 1. Stress increases neuronal activity in the SuM

c-Fos protein expression was assessed after acute stress exposure to test whether the SuM was activated (Figure 1 A). The number of c-Fos^+^ cells was significantly increased by foot shock exposure (Figure 1 B-C). To investigate if SuM would be responsive to diverse stressors, we next examined whether chronic stress, which different mechanism underlying, affects neuronal activity in the SuM (Figure 1 D-E). We chose in vivo electrophysiological extracellular recordings to reveal the neuronal activity before and after chronic social defeat stress, the data shows that the firing rate of regular-spiking neurons (RNs) (Figure 1 F) but not fast-spiking neurons (FNs) (Supplemental Figure 1 A-B) increased after CSDS. Regarding local field potentials, there were no noticeable difference between the naïve and CSDS groups according to power spectrum analysis (Supplemental Figure 1 C-D). These results indicate that acute and chronic stress can strongly activate the supramammillary nucleus.

**Figure 1.**
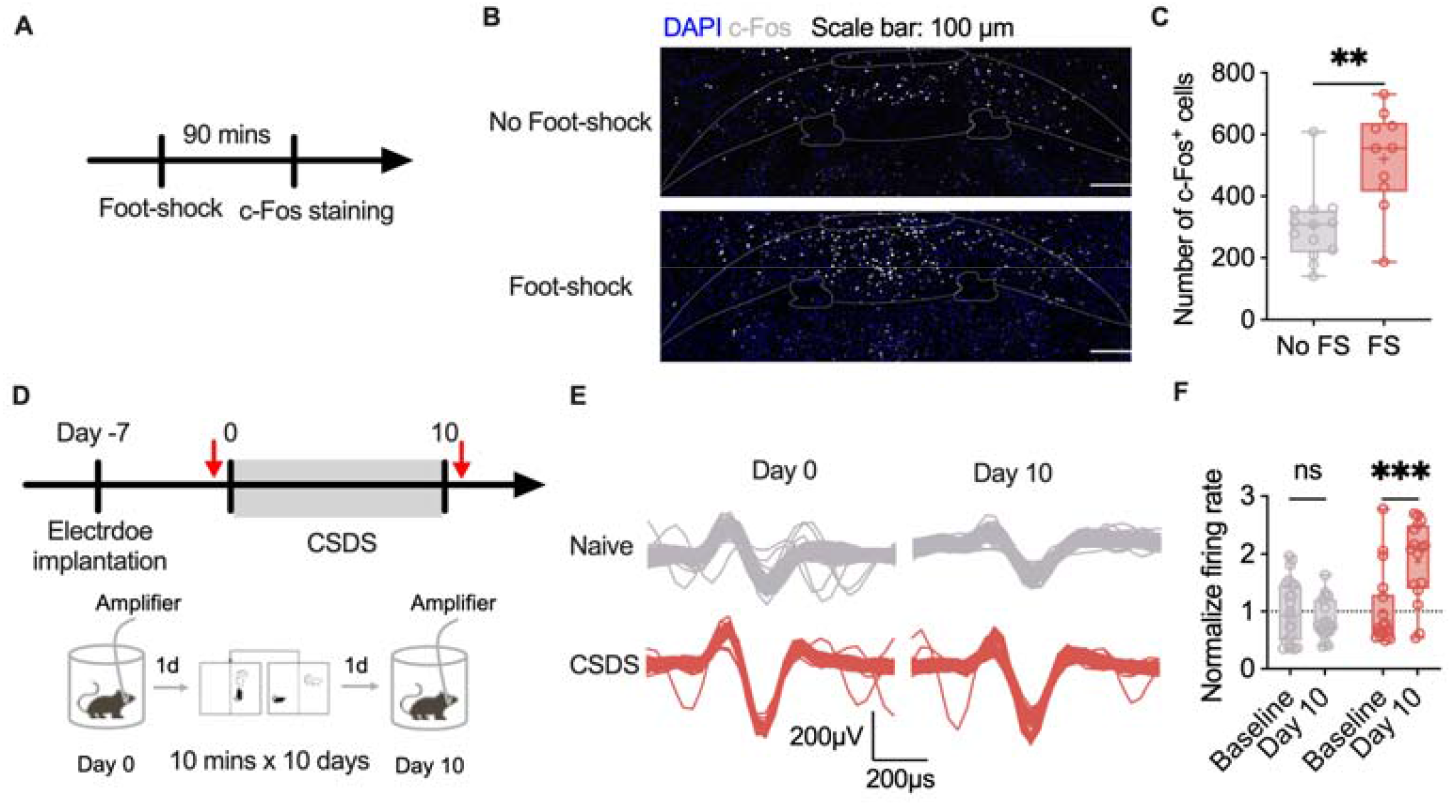
Stress activates the SuM. **(A)** Workflow of the c-Fos staining. **(B)** Representative images of c-Fos staining (DAPI: blue; c-Fos: white; scale bar: 100 µm). **(C)** Statistical analysis of the number of c-Fos-positive cells displayed in panel B. n = 10–13 per group; unpaired t test. **(D)** Workflow of CSDS exposure and in vivo recording. **(E)** Representative spikes acquired by multichannel recording. **(F)** Statistical analysis of the firing rates of RNs at baseline and after CSDS exposure. n = 16□23 per group; two-way ANOVA followed by Sidak’s post hoc test. The bars in C and F indicate the Min to Max of all data point, and the “+” indicates mean value of all data point. “**”, *p* <0.01; “***”, *p* < 0.001. CSDS: chronic social stress; FS: foot shock. Also see **Supplemental Figure 1**.

### 2. Activation of SuM produces anxiety-like behavior

After confirming the activation of SuM caused different types of stress, we then further investigate whether the SuM regulates anxiety behavior in mice. Chemogenetic manipulations were conducted to activate SuM neurons. The experiments were performed as shown in the workflow (Figure 2 A-B). The mice were subjected to the open field (OF) and EZM tests at least 2 weeks after virus injection, followed by a reward-seeking test (Figure 2 E-H, Supplemental Figure 2 A). CNO was administered intraperitoneally 30 minutes before the test. Chemogenetic activation of the SuM did not affect the performance of mice in the OF test (Figure 2 E-F, Supplemental Figure 2 B). Compared with control mice, mice in which the SuM was activated explored the open arms of the EZM less (Figure 2 G), despite no change in distance traveled (Supplemental Figure 2 C). Moreover, mice in which the SuM was activated consumed less food than control mice did (Figure 2 F). These data suggest that there are neuronal ensembles that control the expression of anxiety behavior.

**Figure 2.**
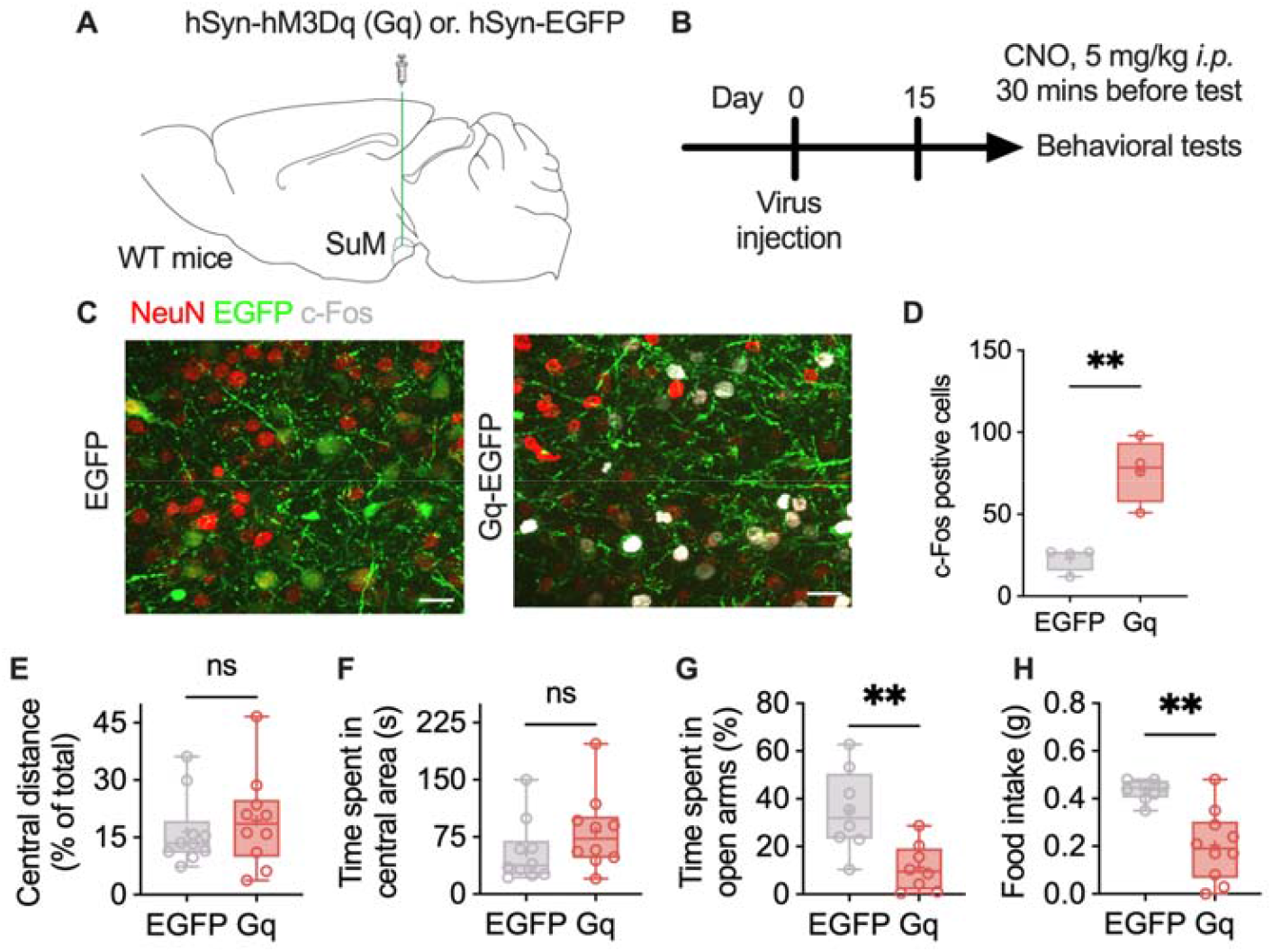
Activation of SuM produces anxiety-like behavior. **(A-B)** Virus injection information (A) and workflow for chemogenetic manipulation (B). **(C)** Representative c-Fos images. **(D)** Statistical analysis of the number of c-Fos-positive cells displayed in panel C. n = 4 per group; unpaired t test. **(E-F)** Statistical analysis of the distance traveled in the central area (E) and the time that the mice spent in the central area (F) in the OF test. n = 10 per group; unpaired t test for the data in (E) and the Mann□Whitney test for the data in (F). **(G)** Statistical analysis of the time that the mice spent in the open arms of the EZM. n = 8 per group; unpaired t test. **(H)** Statistical analysis of sucrose pellets consumed. n = 8–10 per group; Mann□Whitney test. The bars in D-H indicate the Min to Max of all data point, and the “+” indicates mean value of all data point. “ns”, *p* >0.05; “**”, *p* <0.01. Also see **Supplemental Figure 2**.

### 3. Identification of stress-activated neurons in the SuM

We next investigated whether an ensemble that encodes stress and controls the expression of anxiety exists. By crossbreeding Fos 2A-iCreERT2(TRAP2) and Rosa26-LSL-tdTomato (Ai14) mice, we generated TRAP2;Ai14 mice in which activated cells were genetically tagged for visualization (Figure 3 A). Foot shock exposure strongly activated neurons in the SuM but not adjacent areas (Figure 3 B-C). We hypothesized that these stress-activated neurons (SANs) respond exclusively to stress, but not to other stimuli such as reward. To validate the activity-dependent labeling in TRAP2;Ai14 mice, we labeled SANs in two cohorts of mice exposed either to the home cage condition or to foot shock. Several days after labeling, the mice were exposed to either sucrose pellets or social stress to induce reward-related or stress-related c-Fos expression, respectively. The reactivation of SANs under reward stimulation and stress was then compared (Figure 3 D-H). Foot shocks dramatically activated and labeled neurons in the SuM (Figure 3 F). Social stress but not reward (presentation of sucrose pellets) induced neuronal activation (Figure 3 G) and led to a much greater chance of reactivation of SANs (Figure 3 H). These data suggest the specific regulatory effect of the SuM on the effects of stress but not reward.

**Figure 3.**
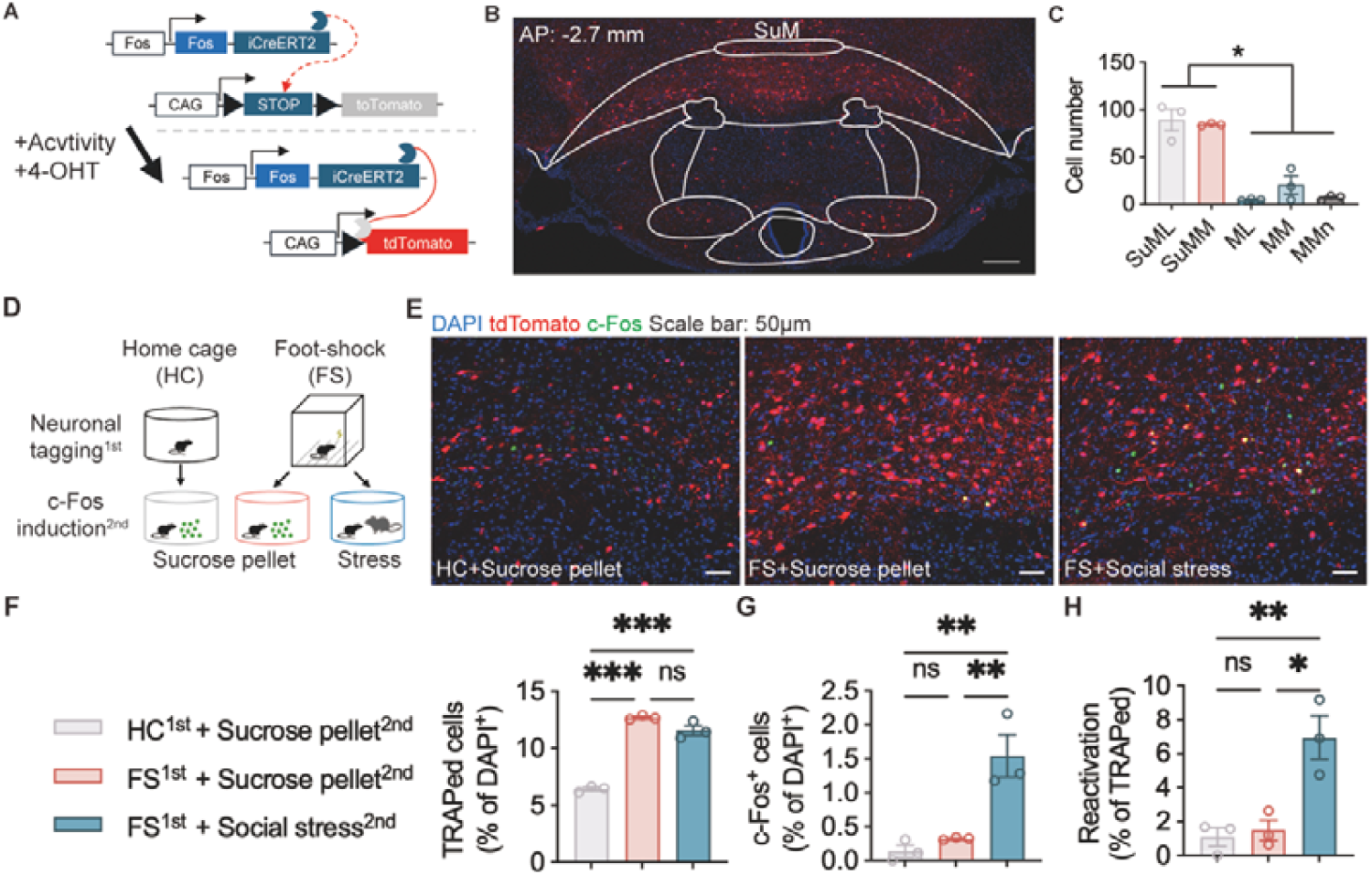
SANs in the SuM selectively respond to social stress but not reward. **(A)** Workflow of neuronal tagging. **(B)** Representative image of SANs in the SuM (DAPI: blue, tagged cells: red). **(C)** Quantitative statistics of stress-tagged cells in several brain areas. n = 3 per area; one-way ANOVA followed by Tukey’4s post hoc test. **(D)** Workflow of neuronal tagging and c-Fos staining. **(E)** Representative images of stress-tagged cells and c-Fos expression induced by sucrose and social stress (DAPI: blue, tdTomato: red, c-Fos: green). **(F)** Statistical analysis of the number of stress-tagged cells in the SuM. n = 3 per group; one-way ANOVA followed by Tukey’s post hoc test. **(G)** Statistical analysis of the number of c-Fos^+^ cells after sucrose or social stress exposure. n = 3 per group; one-way ANOVA followed by Tukey’s post hoc test. **(H)** Statistical analysis of the reactivation of stress-tagged cells. n = 3 per group; one-way ANOVA followed by Tukey’s post hoc test. The data in C and F-H are presented as the means ± SEMs. “ns”, *p* >0.05; “*”, *p* <0.05; “**”, *p* <0.01; “***”, *p* < 0.001.

### 4. Reactivation of SuM^SANs^ promotes anxiety-like behavior

To make sure mice are on similar basal condition while applying chemo-genetic manipulation, we subjected mice to an acute stress protocol involving foot shocks and then performed the elevated plus maze (EPM) and elevated zero maze (EZM) tests to evaluate anxiety on days 2 and 7 (Figure 4 A). The mice that experienced foot shocks showed decreases in the exploration time in the open arms on day 2. However, acute stress-induced anxiety was not detected on day 7 (Figure 4 B), which allow us to compare the reactivation of SANs produced anxiety-like behavior between groups at the same baseline. Seven days after SANs tagging, specific activation of SANs significantly increased the concentration of corticosterone (a peripheral indicator of stress) in the mouse serum (Figure 4 C-D). Experiments involving chemogenetic manipulation also revealed the sound-selective activation of SANs in the SuM (Figure 4 E-F). We then tested whether manipulating SANs in the SuM influences the anxiety-like behavior of the mice. The mice were subjected to the OF and EZM tests at least 1 week after SANs were tagged, followed by reward-seeking tests (Figure 4 G-H). CNO was administered intraperitoneally 30 minutes before the test. Chemogenetic activation of SANs decreased the total distance traveled by the mice in the OF and EZM tests (Supplemental Figure 2 F-G). The mice also presented decreases in the distance traveled in the central area (Figure 4 I) and time spent in the central area in the OF test (Figure 4 J), time spent in the open arms in the EZM test (Figure 4 K) and food consumption (Figure 4 L). These data suggest that SANs in the SuM encode anxiety-like behavior.

**Figure 4.**
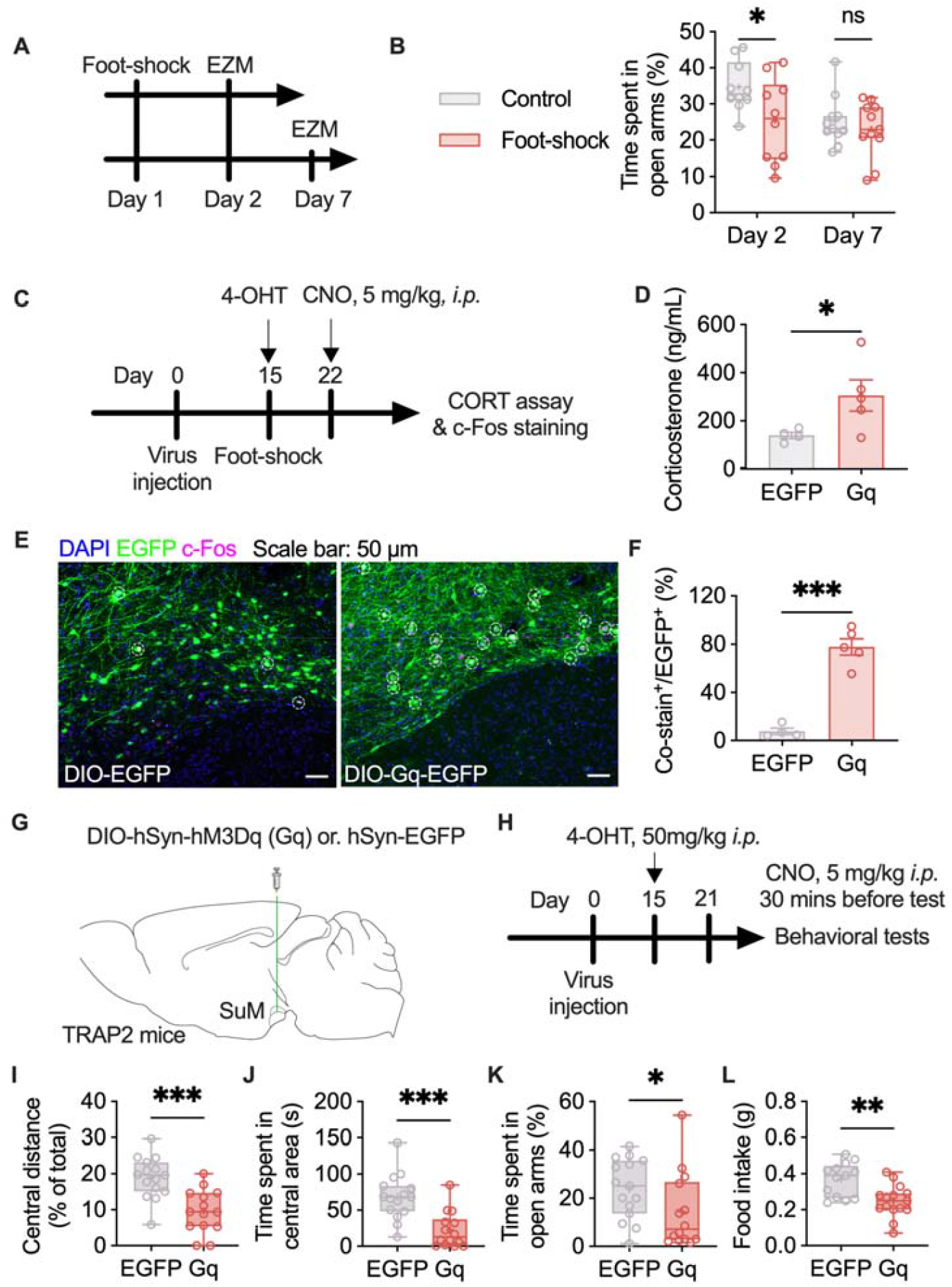
Selective chemogenetic activation of SANs elevates corticosterone level and produces anxiety-like behavior. **(A)** Workflow of the acute stress and anxiety tests. **(B)** Statistical analysis of the time that the mice spent in the open arms of the EZM. n = 10□11 per group; two-way ANOVA followed by Sidak post hoc test. **(C)** Workflow of the CORT assay and c-Fos staining. **(D)** Statistical analysis of the serum concentration of corticosterone after the application of CNO. n = 4-5 per group; unpaired t test. **(E)** Representative images of stress-tagged cells and c-Fos expression induced by chemogenetic manipulation (DAPI: blue, EGFP: green, c-Fos: violet). **(F)** Statistical analysis of the percentage of costained cells relative to EGPF^+^ cells in the SuM. n = 4-5 per group; unpaired t test. **(G-H)** Virus injection information and workflow of chemogenetic manipulation. **(I-J)** Statistical analysis of the distance traveled in the central area (I) and the time that the mice spent in the central area (J) in the OF test. n = 14–15 per group; unpaired t test. **(K)** Statistical analysis of the time that the mice spent in the open arms of the EZM. n = 14–15 per group; Mann□Whitney test. **(L)** Statistical analysis of sucrose pellets consumed. n = 13–15 per group; unpaired t test. The data in D & F are presented as the means ± SEMs. The bars in B and I-L indicate the Min to Max of all data point, and the “+” indicates mean value of all data point. “*”, *p* <0.05; “**”, *p* <0.01; “***”, *p* < 0.001. Also see **Supplemental Figure 2**.

### 5. vSub-SuM projections encode anxiety-like behavior

The SuM receives afferents from various brain areas, and we identified projections to the SuM by using a nonvirus- and virus-based retrograde tracing strategy (Figure 5 A, Supplemental Figure 3 A-C). Afferents from the dSub and vSub were identified using CTB-647 and AAV (Supplemental Figure 4 A-B). These projection neurons expressed Vglut1 RNA but not Vgat RNA (Figure 5 B), suggesting that Sub-SuM projections are excitatory neuronal projections, as Vglut1 is a crucial marker of glutamatergic neurons. We then performed an electrophysiological experiment. Optogenetics-evoked postsynaptic currents in SuM neurons were blocked by perfusion with DNQX, which indicates the existence of glutamatergic projections from the Sub to the SuM (Figure 5 C-E). To investigate how Sub-SuM projections modulate stress and anxiety-like behavior, we then used fiber photometry to measure the calcium concentration to assess the activity patterns of the projection neurons (Figure 5 F-H). The projection neurons in the vSub but not those in the dSub were more strongly activated when the mice moved into the open arms from the closed arms of the EZM, (Figure 5 I-M). On the other hand, vSub but not dSub projection neurons show lower calcium activity while mice backing to the closed arms (Supplemental Figure 5 A-D). Following exposure to acute stress, both dSub-SuM and vSub-SuM projection neurons presented increased calcium activity (Figure 5 N-R). These data suggest that vSub-SuM projections, but not dSub-SuM projections, may participate in regulating anxiety-liker behavior.

**Figure 5.**
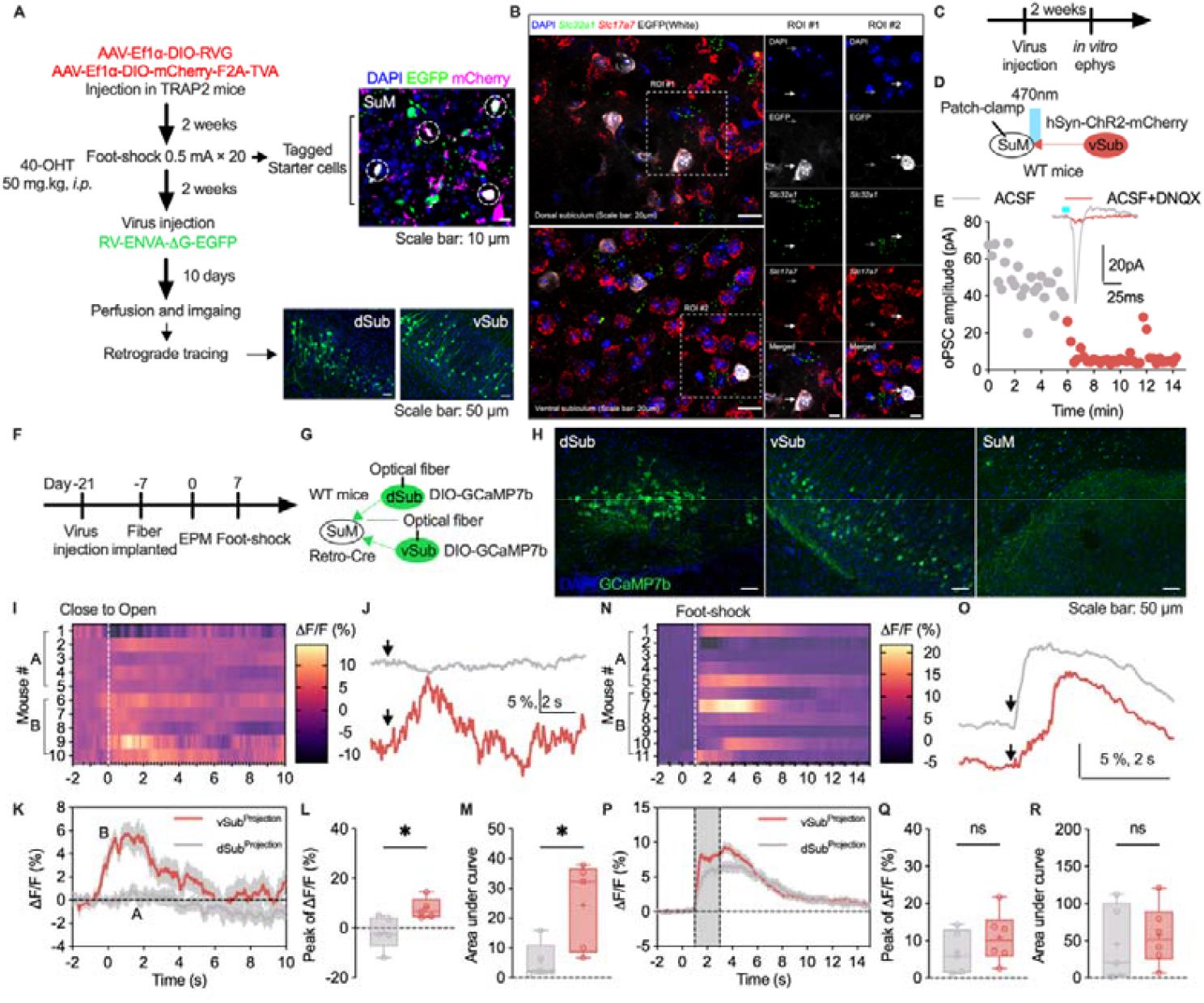
vSub-SuM projections encoding anxiety-like behavior. **(A)** Workflow of virus-based retrograde neuronal tracing. **(B)** Representative images of in situ RNA staining (DAPI: blue, Slc32a1: green, Slc17a7: red, EGFP: white). **(C)** Workflow of ex vivo electrophysiological recording. **(D)** Schematic of oPSCs in the SuM. **(E)** Representative traces of oPSCs. **(F)** Workflow of Ca^2+^ imaging. **(G)** Schematic of Ca^2+^ imaging of dSub and vSub projection neurons. **(H)** Representative images of GCaMP7b expression in the dSub, vSub and SuM (DAPI: blue, GCaMP7b: green). **(I)** Heatmap of the Ca^2+^ fluorescence intensity during the transition from the closed to the open arms. **(J)** Representative Ca^2+^ activity during the transition from the closed arms to the open arms. **(K)** Average ΔF/F of Ca^2+^ recorded in the dSub and vSub. **(L)** Statistical analysis of the peak Ca^2+^ activity. n = 5 per group; unpaired t test. **(M)** Statistical analysis of the area under the curve of Ca^2+^ activity. n = 5 per group; unpaired t test. **(N)** Heatmap of the Ca^2+^ fluorescence intensity during exposure to foot shocks. **(O)** Representative Ca^2+^ activity during exposure to foot shocks. **(P)** Average ΔF/F of Ca^2+^ recorded in the dSub and vSub. **(Q)** Statistical analysis of the peak Ca^2+^ activity. n = 5□6 per group; unpaired t test. **(R)** Statistical analysis of the area under the curve of Ca^2+^ activity. n = 5□6 per group; unpaired t test. The bars in L-M and Q-R indicate the Min to Max of all data point, and the “+” indicates mean value of all data point. “ns”, *p* >0.05; “*”, *p* <0.05. Also see **Supplemental Figure 4 & Supplemental Figure 5**.

### 6. Chronic inhibition of vSub-SuM projections alleviates anxiety-like behavior

After confirming the regulatory role of vSub-SuM projections in anxiety, we hypothesis that inhibition of this projection would alleviate chronic stress-induced anxiety. In the following experiments, the activity of these projections was chronically inhibited via a chemogenetic strategy. The mice were exposed to CSDS after the expression of the Gi protein was induced specifically on vSub-SuM projection neurons and their axons (Figure 6 A-B, D). The body weights of the mice were monitored throughout the entire procedure to assess their health (Figure 6 C). The mice showed no change in social interaction test scores after CSDS exposure (Figure 6 E). In the EZM test, the mice in which vSub-SuM projections were inhibited presented less anxiety-like behavior, as indicated by a longer time spent in the open arms of the EZM but no significant change in the distance traveled (Figure 6 F-G). Taken together, these data suggest that vSub-SuM projections are the essential neuronal projections for regulating chronic stress induced anxiety-like behavior.

**Figure 6.**
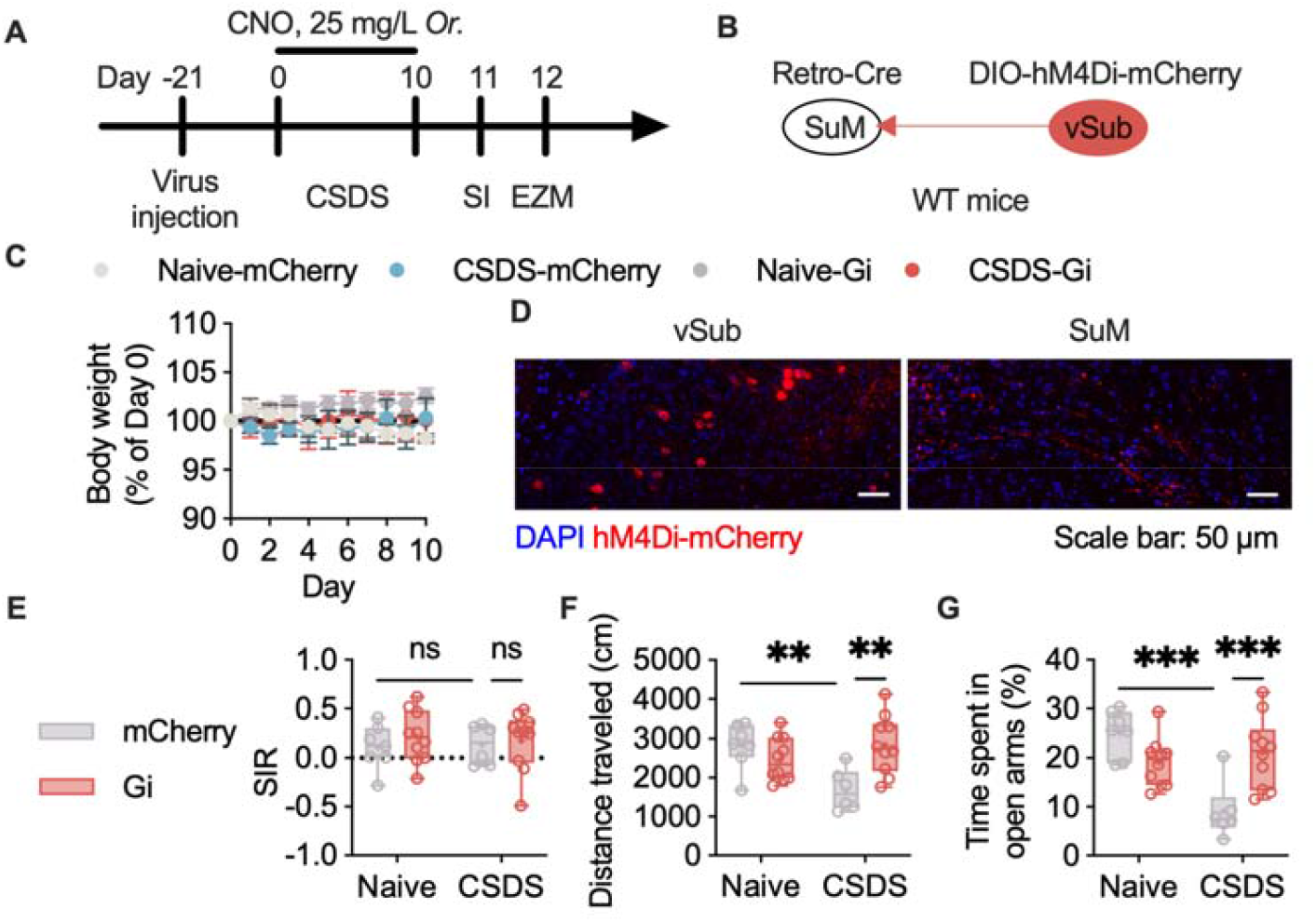
Selective inhibition of vSub-SuM projections alleviated CSDS-induced anxiety. **(A)** Workflow of CSDS and chemogenetic manipulation. **(B)** Schematic of chemogenetic manipulation of specific projections. **(C)** Body weight during CSDS exposure. **(D)** Representative images of virus expression. **(E)** Statistical analysis of the social interaction ratio after CSDS exposure. n = 6-10 per group; two-way ANOVA followed by Sidak’s post hoc test. **(F)** Statistical analysis of the distance that the mice traveled in the EZM. n = 6-10 per group; two-way ANOVA followed by Sidak’s post hoc test. **(G)** Statistical analysis of the time that the mice spent in the open arms of the EZM; two-way ANOVA followed by Sidak post hoc test. The bars in E-G indicate the Min to Max of all data point, and the “+” indicates mean value of all data point. “ns”, *p* >0.05; “*”, *p* <0.05; “**”, *p* <0.01; “***”, *p* < 0.001.

## Discussion

In this study, we combined multiple methods to determine whether the SuM is a brain region that involve in modulating anxiety. SuM neurons strongly respond to acute and chronic stress, and their activation results in robust increases in anxiety-like behavior in mice. We then defined a small ensemble of neurons that are activated by stress, called SANs. These SANs specifically respond to stressful stimuli but not reward. Selective activation of SANs in the SuM increases the serum concentration of corticosterone and anxiety-like behavior in mice. The neuronal circuits that may be underlie the regulation of anxiety were also determined in this study. The subiculum sends glutamatergic projections to the SuM and can be activated by stress, whereas only the vSub has a potential effect on the transition to anxiety. We finally determined that inhibition of SANs in the vSub project to the SuM is sufficient to alleviate anxiety in mice after CSDS exposure.

### Regulation of anxiety avoidance by the SuM

The SuM has been demonstrated to respond to novel environments, social stimulation[7, 33], and stress[34, 35]. Given these findings, we assume that the SuM may be activated by foot shocks, a quantifiable acute stressor used in animal studies. Consistent with this hypothesis, we found that the SuM robustly positively responds to both acute and chronic stress through observable increases in c-Fos expression and increases in the neuronal firing rate. The activation of the SuM was also demonstrated to be essential for maintaining arousal[6, 13]. Sensitization to stressful events and high arousal are often associated with anxiety[36]. Thus, our data strongly suggest that the SuM potentially modulates anxiety. To further confirm whether the SuM participates in anxiety regulation, we recorded neuronal action potentials via multichannel extracellular recording while the mice were moving in the EPM, a traditional type of maze used to test anxiety in rodents. The change in the neuronal firing frequency when mice transitioned from the open arms to the closed arms supports the idea that the SuM may somewhat modulate anxiety. We then manipulated neuronal activity in the SuM via a chemogenetic method and subjected the mice to the EZM test, an improved test for assessing anxiety in rodents. The decrease in the exploration time in the open arms by mice in which the SuM was activated by hM3Dq indicated increased anxiety. We noted that these results are inconsistent with those of some previous reports. Some studies have reported that lesions in the SuM and adjacent areas decrease anxiogenic behavior in rats[15, 37, 38]. López-Ferreras et al. performed the OF test, a complicated test, and reported that the chemogenetic activation of neurons in the SuM results in increased anxiety-like behaviors in rats[16, 17]. However, further experiments involving specific test (e.g., the EZM test) are needed to confirm whether there is a potential difference across species.

### The role of stress-activated neurons in regulating anxiety avoidance

Recent studies have highlighted the importance of activity-tagged neuronal ensembles in regulating various behaviors, particularly memory[20, 39–41], food consumption[22], the inflammatory response[24] and emotion[26, 42]. A negative experience-related neuronal ensemble in the hippocampus was found to increase susceptibility to chronic stress[26]. A recent study reported that the lateral habenula contains a small population of neurons that are recruited in response to stress and mediate the development of depression in mice[42]. These studies suggest that SANs may be important for emotional regulation. In this study, we found that the SuM was more strongly activated by acute stress than was adjacent areas. SANs were more likely to be reactivated by social stress than by sucrose reward, indicating their potential to specifically encode anxiety. The serum corticosterone concentration can be used as a marker of stress-induced change in the peripheral blood. Previous studies showed serum corticosterone can be increased by various stress stimulation[39–42]Click or tap here to enter text.; meanwhile, intentionally supplementing the diet with corticosterone can induce anxiety-like behaviors in rodents[43]. Our data showed that the chemogenetic activation of SANs in the SuM increased the serum corticosterone concentration, whereas the inactivation of SANs had no effect, suggesting the chemogenetic manipulation of SANs may cause similar anxiety effect like real stressors. These findings in combination with the results of the OF and EZM suggest that SANs in the SuM are more likely involved in modulating anxiety-like avoidance. However, the reactivation rate of SANs caused by different stressor was relatively lower than the initial activation rate caused by foot shock (Figure 3). This suggests that stress-activated neuronal clusters may have more flexible recruitment principles, with only a small number of neurons potentially encoding emotional information, while most other neurons remain involved in encoding other neural activities. Studies in other field, particularly studies of memory engram, has shown that the sets of neurons activated during learning are dynamic and exhibit high flexibility[44, 45]. While the activation of SANs produced anxiety-like behavior, the future study will examine whether silencing SuM SANs, either during stress exposure or during anxiety testing, can prevent or reduce stress-induced anxiety. We also found that both the nonselective activation of SuM neurons and the selective activation of SANs in the SuM significantly suppressed the consumption of sucrose pellets. This result may be attributed to the anxiety-induced suppression of reward seeking[43]. However, further experiments are still needed to confirm whether this effect is anxiety dependent and whether basal food consumption is affected.

### A relevant neural circuit that regulates anxiety avoidance

The SuM recruits and is targeted by neuronal projections in the hippocampus, medial septum, and cortex[32]. To further understand the circuitry through which the SuM regulates anxiety, we identified projections from the dSub and the vSub to the SuM[46]. Fiber photometry was used to measure the Ca^2+^ concentration in projection neurons in the dSub and vSub, and the results revealed increased Ca^2+^ activity in vSub-SuM projection neurons but not dSub-SuM projection neurons when the mice transited from the closed arms to the open arms, indicating that vSub-SuM projections encode anxiety. To confirm the regulation of anxiety by vSub-SuM projections, we exposed mice to CSDS and found that constant inhibition of vSub-SuM activity significantly abolished CSDS-induced anxiety in mice. Unlike the dorsal hippocampus, which is involved in the regulation of cognition, the ventral hippocampus is often involved in regulating emotion[47]. The ventral CA1 area and its projections to the lateral hypothalamic area were found to mediate innate anxiety, and its activation increases anxiety-like behavior in mice[28]. Although very close spatially, neurons in the subiculum are somewhat different from those in the CA1 region[48, 49]. The vSub and its downstream brain areas were found to regulate anxiety[50, 51]. Jing-Jing et al. reported that the vSub and its projections to the anterior hypothalamic nucleus are essential for anxiety because the inhibition of these projections decreases anxiety-like behavior[27]. Our data are consistent with these findings and suggest the mediating role of the vSub and its projections to different subareas in the hypothalamus.

In summary, the activation of SuM increases anxiety-like behavior. Stressful event recruits a neuronal ensemble in SuM. The activation of SANs also significantly increases anxiety-like behavior and suppresses reward seeking. SuM receives glutamatergic projections from the vSub, and inhibition of these projections can diminish CSDS-induced anxiety-like behavior. These results suggest that SuM plays important role in regulating anxiety-like behavior and furthermore studies are worthy performing.

## Acknowledgment

We thank Dr. Wenting Wang for his generous gift of the transgenic mice. We thank all the members of MTT for their valuable comments.

This research was funded by the National Natural Science Foundation of China (No. 82371518, 82071516, and 82441060), STI 2030—Major Projects 2021ZD0200500, the Humanities and Social Science Fund of the Ministry of Education of China (No. 22XJC880005), the Innovation Capability Support Program of Shaanxi (Program No. 2021PT-055), the Natural Science Basic Research Plan in Shaanxi Province of China (Program No. 2024JC-YBQN-0902 and 2023-JC-YB-189), and the Scientific and Technological Innovation Team of Shaanxi Innovation Capability Support Plan (No. 2022TD-47).

## Disclosures

The authors declare that they have no competing interests.

## Methods and Materials

### Animals

Male C57BL6/J mice aged 12–20 weeks were used. Fos^2A-iCreERT2^ (TRAP2) mice were a gift from Wenting Wang (JAX, cat. No. 030323). Rosa26-CAG-LSL-tdTomato (Ai14) mice were purchased from the Shanghai Model Organisms Center (cat. No. NM-KI-225042). Male CD-1 mice aged 8–10 months were purchased from Charles River (cat. No. 201). To construct TRAP2;Ai14 mice, homozygous male TRAP2 mice and homozygous female Ai14 mice were bred. Homozygous TRAP2;Ai14 mice were maintained and used in the experiments. We performed genotyping for TRAP2 and Ai14 via PCR with the following primers: TRAP2 (wild-type: 357 bp, mutant: 232 bp): wild-type forward: GTCCGGTTCCTTCTATGCAG, mutant forward: CCTTGCAAAAGTATTACATCACG, common: GAACCTTCGAGGGAAGACG; Ai14 (wild-type: 297 bp, mutant: 196 bp): wild-type forward: AAGGGAGCTGCAGTGGAGTA, wild-type reverse: CCGAAAATCTGTGGGAAGTC, mutant forward: GGCATTAAAGCAGCGTATCC, mutant reverse: CTGTTCCTGTACGGCATGG.

The mice were housed 4–5 per cage at a constant temperature and humidity (22 ± 1°C, 30–40% RH)on a day–night cycle (lights on from 08:00–20:00) with a fixed-intensity light source. Each mouse was acclimated to the testing environment for 1□2 min. Acclimation was performed for three days before the behavioral experiments. The mice were fed ad libitum and euthanized with CO_2_ after all the tests were finished. The experimental protocols described here were approved by the Animal Ethics Committee of Shaanxi Normal University.

### Behavioral procedure

### Acute stress exposure

The mice were exposed to acute stress according to a previously reported procedure[52]; specifically, they were exposed to 20 foot shocks with an intensity of 0.5 mA that were randomly delivered across 10 minutes. The foot shocks were delivered in a fear conditioning box (Med-associates).

### Chronic stress exposure

We used a CSDS protocol to induce anxiety and depression in the mice[53]. Each C57BL6/J mouse was housed with one CD-1 mouse, and the mice were separated by a transparent plexiglass board with several small holes. The mice were allowed to contact each other directly for 10 minutes every day for 10 days. Body weight was measured and recorded every day before contact.

### OF test

The OF test was carried out in a 50 × 50 × 35 cm arena made of white plexiglass. The mice were allowed to move freely in the arena for 10 minutes, and the distance the mice traveled and the time the mice spent in the central area were recorded and analyzed.

### EPM test

The EPM consisted of two open arms (30 × 7 cm), two closed arms (30 × 7 × 14 cm) and a central area (7 × 7 cm). The mice were allowed to move freely in the arena for 10 min, and the time the mice spent in the open arms was recorded and analyzed.

### EZM test

The EZM was used to test whether the mice were anxious. The EZM used in this study was made of organic glass (height of 60 cm), with an inner diameter of 51.8 cm and an outer diameter of 65 cm. The closed arms of the EZM were separated by two 15 cm-high pieces of organic glass, the outer one of which was opaque. After a 15-minute habituation period, the mice were placed into the EZM and allowed to move freely for 10 minutes. Videos were recorded and analyzed using EthoVisionXT software. The time that the mice spent in the open arms was compared between the groups to evaluate anxiety-like behavior.

### Reward seeking

On the first day, sucrose pellets were provided for habituation. The mice were then deprived of food on the second day. On the third day, the mice were placed in a new home cage without bedding and allowed to eat freely for 2 hours. The pellets were weighed to evaluate whether the experimental manipulation influenced reward seeking by the mice.

### Social interaction test (SIT)

The SIT was conducted as previously described[53]. The SIT involved two 2.5-minute phases. In the first phase (no-target phase), we placed each C57BL6/J mouse in the periphery of the arena opposite the social interaction area (SIA). We allowed the animal to explore the arena freely. In the second phase (with-target phase), each C57BL6/J mouse was placed in the arena again, with a new CD-1 mouse in the SIA. The social interaction ratio (SIR) was calculated using the following formula:

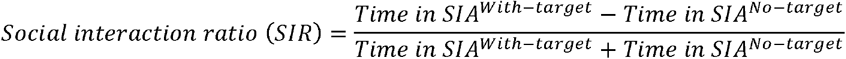

Click or tap here to enter text.**Neuronal tagging of stress-activated neurons**

To specifically label SANs, TRAP2 or TRAP2;Ai14 mice were intraperitoneally (i.p.) injected with 4-hydroxytamoxifen (4-OHT, 50 mg/kg) immediately after acute stress exposure or the learning phase of the CFC test. The mice were subjected to the next experiment or test after 7 days to allow Cre-dependent recombination.

4-OHT (CAS No. 68392-35-8. Sigma, cat. No. H6278 or Bidepharm, cat. No. BD00958757) was dissolved in DMSO at a concentration of 62.5 mg/mL and diluted with vehicle (containing 10% TWEEN-80 and 80% saline) on the day of neuronal tagging. The final concentration of DMSO was kept below 10% to avoid toxicity.

### Observation of the reactivation of SANs

One week after neuronal tagging, whether previously tagged SANs were reactivated in TRAP2;Ai14 mice when they were subjected to social stress was assessed. The mice in the first group were underwent neuronal tagging in their home cages, and c-Fos expression was induced by sucrose pellets. The mice in the second group were subjected to neuronal tagging in response to foot shock exposure, and c-Fos expression was induced by sucrose pellets. The mice in the third group were subjected to neuronal tagging in response to foot shock exposure, and c-Fos expression was induced by social stress (one CD-1 mouse was placed in the home cage). The mice were then sacrificed 90 minutes after sucrose pellet feeding or social stress exposure, and c-Fos immunofluorescence staining was performed. The number of c-Fos-positive neurons was counted to determine whether reward and cross-strain social stress could activate neurons in the SuM. The likelihood of SAN reactivation was calculated using the following formula:

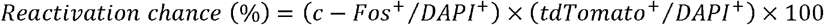

### Viral vectors

An adeno-associated virus (AAV) vector was used to label and manipulate specific neurons or determine the calcium concentration. To manipulate the neuronal activity in the SuM, AAV2/9-hSyn-hM3Dq-EGFP (titer: 5.00E+12 GC/mL, BrainVTA, cat. No. PT-0891) or its control vector AAV2/9-hSyn-EGFP (titer: 5.00E+12 GC/mL, BrainVTA, cat. No. PT-1990) was injected into the SuM of mice.

To manipulate the activity of SANs in the SuM, AAV2/9-hSyn-DIO-hM3Dq -EGFP (titer: 5.00E+12 GC/mL, BrainVTA, cat. No. PT-0891) or its control vector AAV2/9-hSyn-DIO-EGFP (titer: 5.00E+12 GC/mL, BrainVTA, cat. No. PT-1103) was injected into the SuM of TRAP2 mice.

To chronically inhibit vSub-SuM circuitry activity, AAV2/Retro-hSyn-Cre (titer: 2.00E+12 GC/mL, Taitool, cat. No. S0278) was injected into the SuM, and AAV2/9-hSyn-DIO-hM4Di-mCherry (titer: 2.00E+12 GC/mL, Taitool, cat. No. S0193) or its control vector AAV2/9-hSyn-DIO-mCherry (titer: 2.00E+12 GC/mL, Braincase, cat. No. BC-0025) was injected into the vSub of wild-type mice.

To determine the calcium concentration in dSub/vSub-SuM projection neurons, AAV2/Retro-hSyn-Cre (titer: 2.00E+12 GC/mL, Taitool, cat. No. S0278) was injected into the SuM, and AAV2/9-hSyn-DIO-GCaMP7b (titer: 5.00E+12 GC/mL, BrainVTA, cat. No. PT-2892) was injected into the dSub/vSub of wild-type mice.

For the ex vivo electrophysiological experiment, AAV2/9-hSyn-ChR2-mCherry (titer: 5.00E+12 GC/mL, BrainVTA, cat. No. PT-0150) was injected into the vSub of wild-type mice.

### Neuronal tracing

Retrograde neuronal tracing was initially performed via injection of a serotype-2 AAV vector (AAV2/Retro-hSyn-EGFP, titer: 5.00E+12 GC/mL, Taitool, cat. No. S0237) and CTB-647 (1 µg/µL, Thermo Fisher, cat. No. C34778) into the SuM of wild-type mice. The mice were then sacrificed after 2 weeks, and the brains were cut into coronal slices for imaging.

To precisely trace neuronal afferents projecting to SuM^SANs^, AAV2/9-Ef1α-DIO-RVG (titer: 5.00E+12 GC/mL, BrainVTA, cat. No. PT-0061) and AAV2/9-Ef1α-DIO-mCherry-F2A-TVA (titer: 5.00E+12 GC/mL, BrainVTA, cat. No. PT-0207) were injected into the SuM of TRAP2 mice simultaneously. The rabies virus (RV) vector RV-ENVA-ΔG-EGFP (titer: 2.00E+08 IFU/mL, BrainVTA, cat. No. R01001) was injected into the SuM 2 weeks after neuronal tagging. The mice were then sacrificed after 2 weeks, and the brains were cut into coronal slices for imaging.

### Stereotaxic surgery

The mice were anesthetized using isoflurane at a concentration of 1.5∼2.0%. Virus was injected into the SuM (AP: -2.8, ML: 0, DV: -4.5 mm), dSub (AP: -2.8, ML: ±0.7, DV: -1.7 mm) or vSub (AP: -3.5, ML: ±3.0, DV: -4.6 mm) according to the experimental design. If only one type of virus needed to be injected into a single brain area, the final volume was typically 150 nL. Otherwise, the final volume of the virus mixture was 200 nL. The viruses were injected at a rate of 50 nL/min. The syringe was held in place for at least 5 minutes and carefully removed from the brain. The mice were then returned to their home cages, and their health was monitored on the following days. All the mice that underwent surgery were subjected to the subsequent experiment after 2 weeks or more to allow virus expression.

For fiber photometry, ceramic ferrules (outer diameter: 2.5 mm, core diameter: 0.2 mm, NA: 0.50) were inserted into the dSub (AP: -2.8, ML: ±0.7, DV: -1.5 mm) or vSub (AP: -3.5, ML: ±3.0, DV: -4.4 mm) 2 weeks after virus injection under the guidance of a laser (wavelength: 470 nm). Calcium imaging was conducted at least 1 week after ferrule implementation.

### Fiber photometry

Commercially available equipment (Thinker Tech) was used to determine the calcium concentration. The fluorescence signal was activated by a laser at 470 nm, and the signal was transmitted through a low-autofluorescence fiber□optic patch cord and rotary (doric lenses) and collected. The final activation intensity was set to ∼40 µW. The sampling rate was 50 Hz for all the recordings. The mice were habituated to the fiber□optic patch cord for 3 consecutive days before recording. A TTL lasting 0.1 s was delivered by the software to mark the timepoint when the mouse moved from a closed arm to an open arms in the EPM (USB-IO box, Noldus). Continuous data were stored as *.tdms files and analyzed using custom-made software in MATLAB.

### Chemogenetic manipulation

To manipulate neuronal activity in the SuM in wild-type and TRAP2 mice, clozapine N-oxide (CNO, 5 mg/kg; Cayman, cat. No. 25780) was injected i.p. 30 minutes before behavioral tests were performed. For chronic inhibition of circuit activity, CNO was administered orally (25 mg/L).

For acute and chronic experiments, CNO was dissolved in DMSO at a concentration of 10 mg/mL and stored at -20°C or in saline at a concentration of 1 mg/mL and stored at -80°C. The storage solution was diluted with saline to a concentration of 0.75 mg/mL to prepare a working solution for acute manipulation or to a concentration of 25 mg/L to prepare a working solution for chronic inhibition on the day of the experiment.

### Immunofluorescence

The mice were anesthetized with 20% urethane and perfused with PBS or saline. The mouse brain was dissected and immersed in 4% paraformaldehyde (PFA) at 4°C overnight. The PFA solution was then replaced with a 30% sucrose solution. After the brain sank to the bottom, it was embedded in optimal cutting temperature (OCT) compound and frozen in a cryostat (CM1950, Leica). Coronal slices (40 µm) were cut and collected in a 24-well plate. After the residual OCT was removed with PBS, the slices were blocked with 0.3% Triton X-100 and 10% normal donkey serum at room temperature (RT) for 2 hours. The slices were then incubated with diluted primary antibody (rabbit anti-c-Fos, 1:500, Cell Signaling Technology, cat. No. 2250) at 4°C overnight. The next day, the slices were washed and incubated with secondary antibody dilutions (donkey anti-rabbit conjugated to AF647, 1:500, Jackson ImmunoResearch, cat. No. 706-605-148) at RT for 2 hours. After washing, the slices were transferred to slides and mounted with an antifade reagent (Thermo Fisher, cat. No. P36981). Images of the slices were collected using a Zeiss M2 microscope and then analyzed.

### RNA fluorescence in-situ hybridization

The samples were processed as described in the *Immunofluorescence* section. Slices (10 µm thick) were cut and dried at RT for ∼15 minutes and then heated at 37°C for 30 minutes in a hybridization oven. The baked slides were then moved to precooled 4% PFA solution for fixation (∼15 minutes). The slices were dehydrated in 100% ethanol at RT for 5 minutes. The dehydration step was then repeated. The following steps were performed as recommended by the manufacturer (ACDbio, cat. No. 323100). To label vglut1, vglut2 and vgat RNA, the slices were hybridized with Mm-*Slc17a7* (ACDbio, cat. No. 416631-C1), Mm-*Slc17a6* (ACDbio, cat. No. 319171-C1) and Mm-*Slc32a1* (ACDbio, cat. No. 319191-C3), respectively. The samples were then stained with Opal dye.

### Costaining of protein and RNA

After confirming RNA staining, the slices were blocked in 10% normal goat serum for 1 hour. The blocking solution was removed, and the slices were incubated with diluted primary antibody (mouse anti-GFP, 1:500, Thermo Fisher, cat. No. MA5-16256; rabbit anti-tdTomato, 1:500, Oasis BioFarm, cat. No. OB-PRB013) at 4°C overnight. The slides were washed with PBS and incubated with secondary antibody solution (goat anti-rabbit/mouse conjugated to HRP, Proteintech, cat. No. PR30009) at RT for 1 hour (in the dark). After washing, the slices were stained with Opal dye at RT for 30 minutes. The slides were then mounted and imaged.

### Corticosterone assay

Mouse whole blood was collected 90 min after CNO injection (5 mg/kg, i.p.). The samples were subsequently centrifuged at 2000×g for 10 minutes at 4°C after being left to stand at RT for 30–60 minutes. The supernatant was then carefully collected as the serum. Corticosterone levels were then measured using a commercial ELISA kit (Beyotime, cat. No. PC100) according to the manufacturer’s instructions.

### Ex vivo electrophysiology

The mice were anesthetized with urethane and then decapitated. The brain was quickly removed from the skull and immersed in precooled sucrose-based cutting solution (in mM, 225 sucrose, 2.5 KCl, 1.25 NaH_2_PO_4_, 26 NaHCO_3_, 11 D-glucose, 5 L-ascorbic acid, 3 sodium pyruvate, 7 MgSO_4_·7H_2_O, 0.5 CaCl_2_). After being fixed on a metal pallet, the brain was cut into 300-µm slices. The slices were then collected and incubated in artificial cerebrospinal fluid (ACSF) containing (in mM) 122 NaCl, 2.5 KCl, 1.25 NaH_2_PO_4_, 26 NaHCO_3_, 11 D-glucose, 2 MgSO_4_·7H_2_O, and 2 CaCl_2_ equilibrated with 95% O_2_-5% CO_2_ at 28°C for at least 1 hour before recording.

To evoke postsynaptic currents (PSCs) using light, whole-cell recording of global SuM neurons near axons illuminated by ChR2-mCherry injected into the vSub was performed. The final light intensity at the end of the optical fiber was set to ∼5 mW/mm^2^. Then, blue light (470 nm, width: 10 ms, frequency: 0.05 Hz) was used to evoke optically induced PSCs (oPSCs). DNQX (20 µM) was perfused into the ACSF to isolate AMPA-dependent currents from the oPSCs after establishing a 5-minute baseline.

### In vivo electrophysiology

The mice were anesthetized with 2% isoflurane and fixed to a stereotaxic device. A sixteen-channel microwire electrode array (KD-MWA, KedouBC), a 4x4 array of 25 µm NiTi wires spaced 200 µm apart, was slowly inserted into the mouse brain. Four small nails were first inserted into the skull, with a ground wire presoldered onto one of them. The electrode array was left in the SuM (AP: -2.8, ML: 0, DV: -4.55 mm), and then dental cement was used to fix it onto the skull.

The mice were introduced to the recording area at least one week after surgery. During the day, the electrode array attached to the mouse skull was connected to the OpenEphys acquisition board through an Intan head stage. An OpenEphys GUI was used to visualize and save electrical signals. The mice were allowed to move freely inside a home cage-like arena for at least 20 minutes. Only data acquired during the last 5 min were saved and then analyzed via Python-based software and a customized Python script.

Spikes were detected and divided into single units using SpikeInterface (https://github.com/SpikeInterface/spikeinterface). Continuous binary raw data (sampling rate: 30 kHz) were imported and filtered using a bandpass butter filter at a cutoff value of 300 Hz. Movement artifacts were removed by subtracting medians across all channels. The templates were then extracted and fitted using spyking-circus2 inside the SpikeInterface frame. Neurons meeting the following criteria were excluded from the subsequent analysis: (1) spikes with refracting period violations smaller than 1 ms, accounting for more than 2% of total spikes, and (2) a total frequency lower than 0.2 Hz. Neurons with spike frequencies ≥ 10 Hz were considered RNs, whereas those with spike frequencies < 10 Hz were considered FNs, as reported in a previous study[7]. The local field potential was extracted and analyzed using the power spectrum analysis tool in MATLAB.

### Statistical analysis

The data are presented as the means ± SEMs in all the figures in this manuscript. For normally distributed data with equal standard deviations, independent t tests for unpaired data and dependent t tests for paired data were performed in GraphPad software to compare mean values between two groups. Otherwise, the Mann□Whitney test for unpaired data and the Wilcoxon test for paired data were performed instead. One-way ANOVA followed by Tukey’s post hoc test and two-way ANOVA followed by Sidak’s post hoc test were performed to compare mean values among more than three groups. A *p* value less than 0.05 was considered to indicate a statistically significant difference between groups. “*” represents *p* < 0.05, “**” represents *p* < 0.01, and “***” represents *p* < 0.001. Detailed statistical information referring to specific figure can be found in Table 1.

## Notes

### Competing Interest Statement

The authors have declared no competing interest.

### Summary of Updates

Title revised; Discussion revised; supplemental file updated. and other modifications according to the reviewers' suggestion.

