## Supplemental Figure 1 for "A stress-activated neuronal ensemble in the supramammillary nucleus produces anxiety-like behavior in male mice"

**Supplementary information**

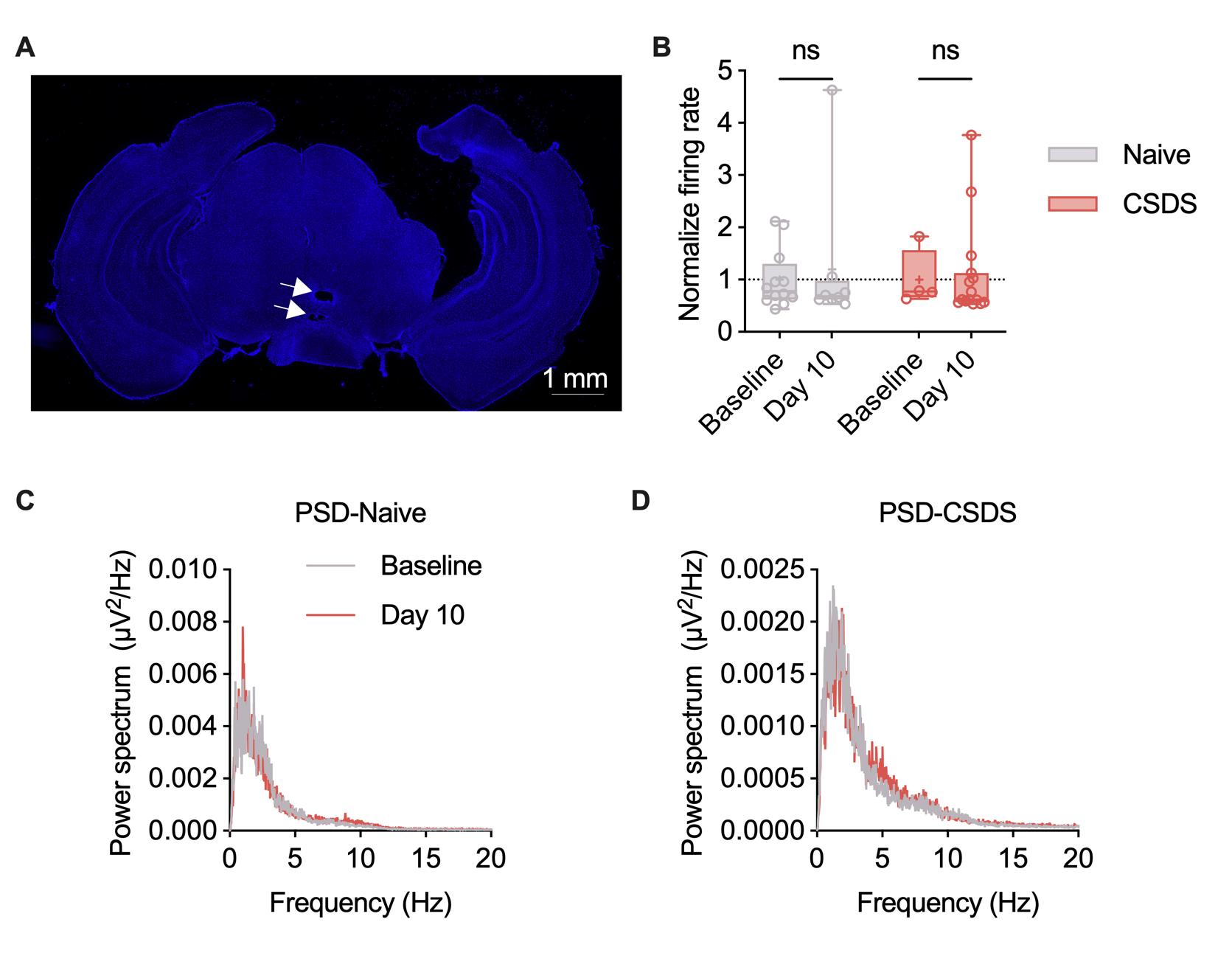

**Supplemental Figure 1.** The effect of CSDS on the LFP in the SuM

**(A)** Representative images of the location of electrodes. **(B)** Statistical comparison of the firing rate of FNs between baseline and after CSDS. n = 4-12 neurons from 2-4 mice per group, two-way ANOVA, Sidak’s post-hoc test. **(C-D)** Power spectrum of local field potentials. Data in B is presented as Mean ± SEM. “ns”*p* >0.05.

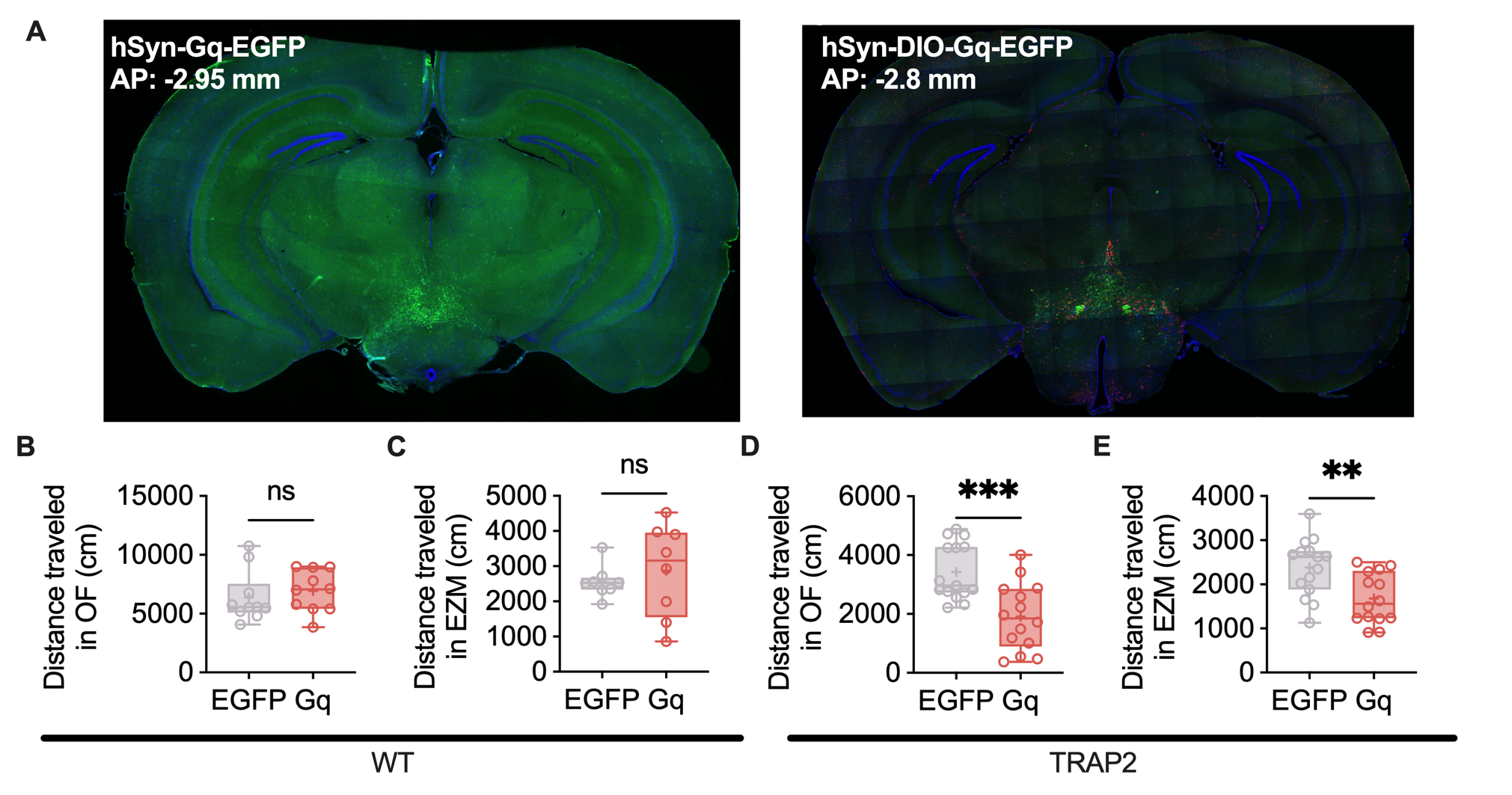

**Supplemental Figure 2.** The chemo-genetic manipulation of SuM and SANs has effects on the performance of mice in the OF and the EZM.

(A) Representative images of virus expression. (B) Statistical comparison of the total distance that WT mice traveled in OF. (C) Statistical comparison of the total distance that WT mice traveled in EZM. (D) Statistical comparison of the total distance that TRAP2 mice traveled in OF. (E) Statistical comparison of the total distance that TRAP2 mice traveled in EZM. Data in B-I are presented as Mean ± SEM. “ns”*p* >0.05, “**”*p* <0.01, “***”*p* < 0.001.

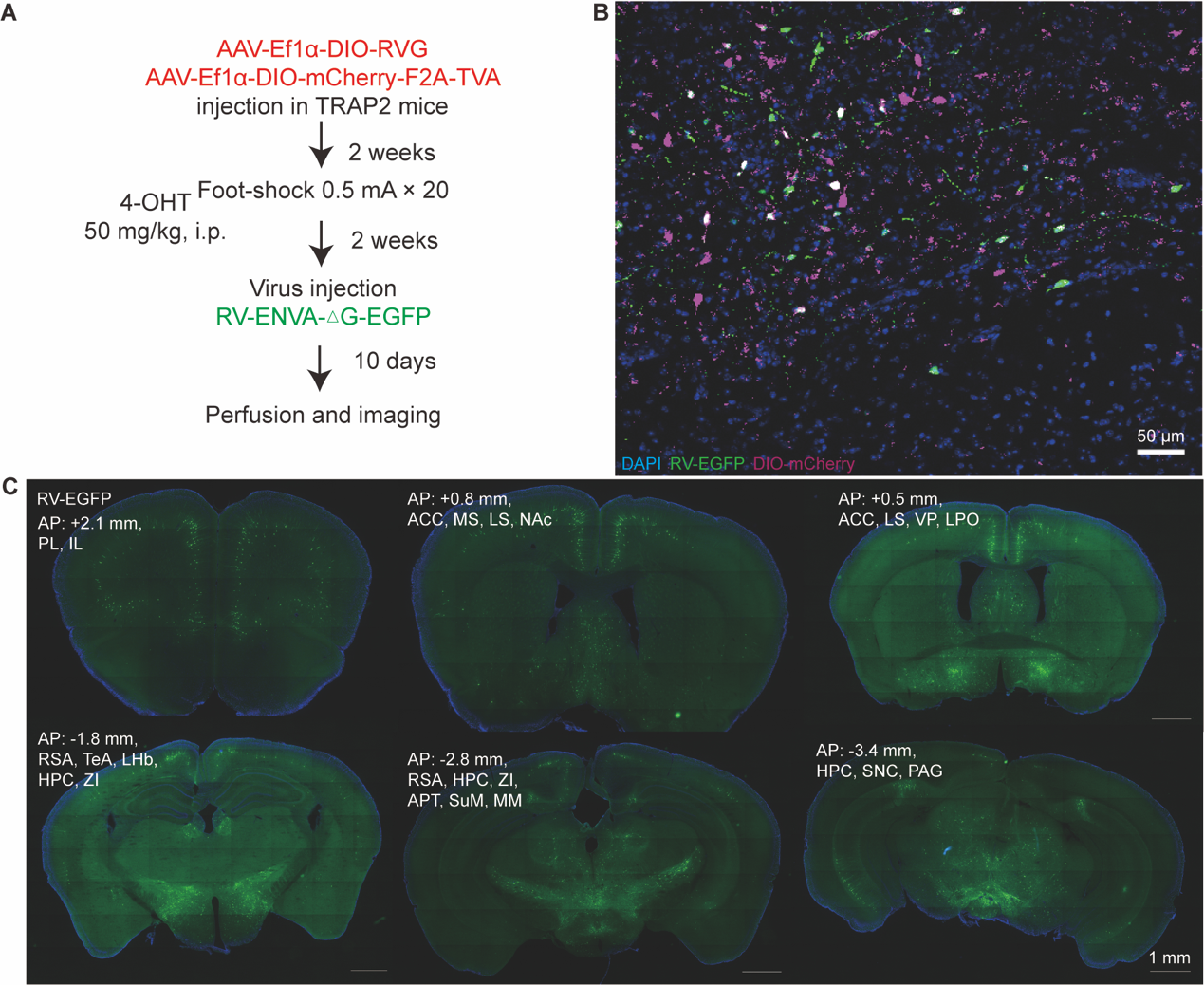

**Supplemental Figure 3.** Specific retrograde neuronal tracing of the upstream of the SAN in the SuM

**(A)** Workflow of RV-based retrograde neuronal tracing. **(B)** Representative image of virus expression in SuM. **(C)** Representative images of traced upstreaming brain area of SuM.

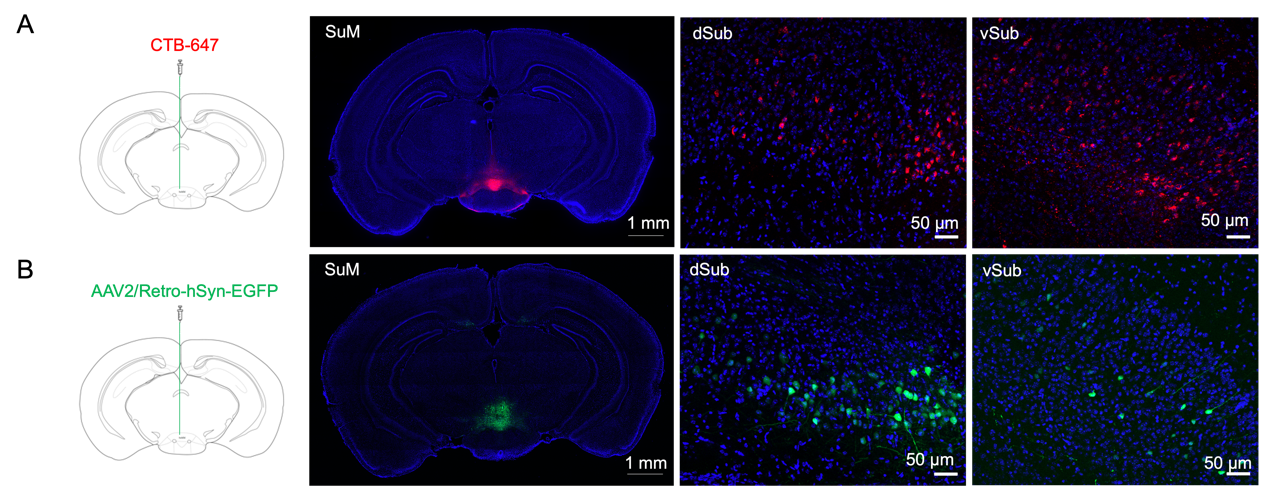

**Supplemental Figure 4.** Non-virus and virus-based retrograde neuronal tracing of the upstream of the SuM

**(A)** Representative images of retrograde neuronal tracing using CTB-647, injection site, dSub and vSub. **(B)** Representative images of retrograde neuronal tracing using AAV2/Retro, injection site, dSub and vSub.

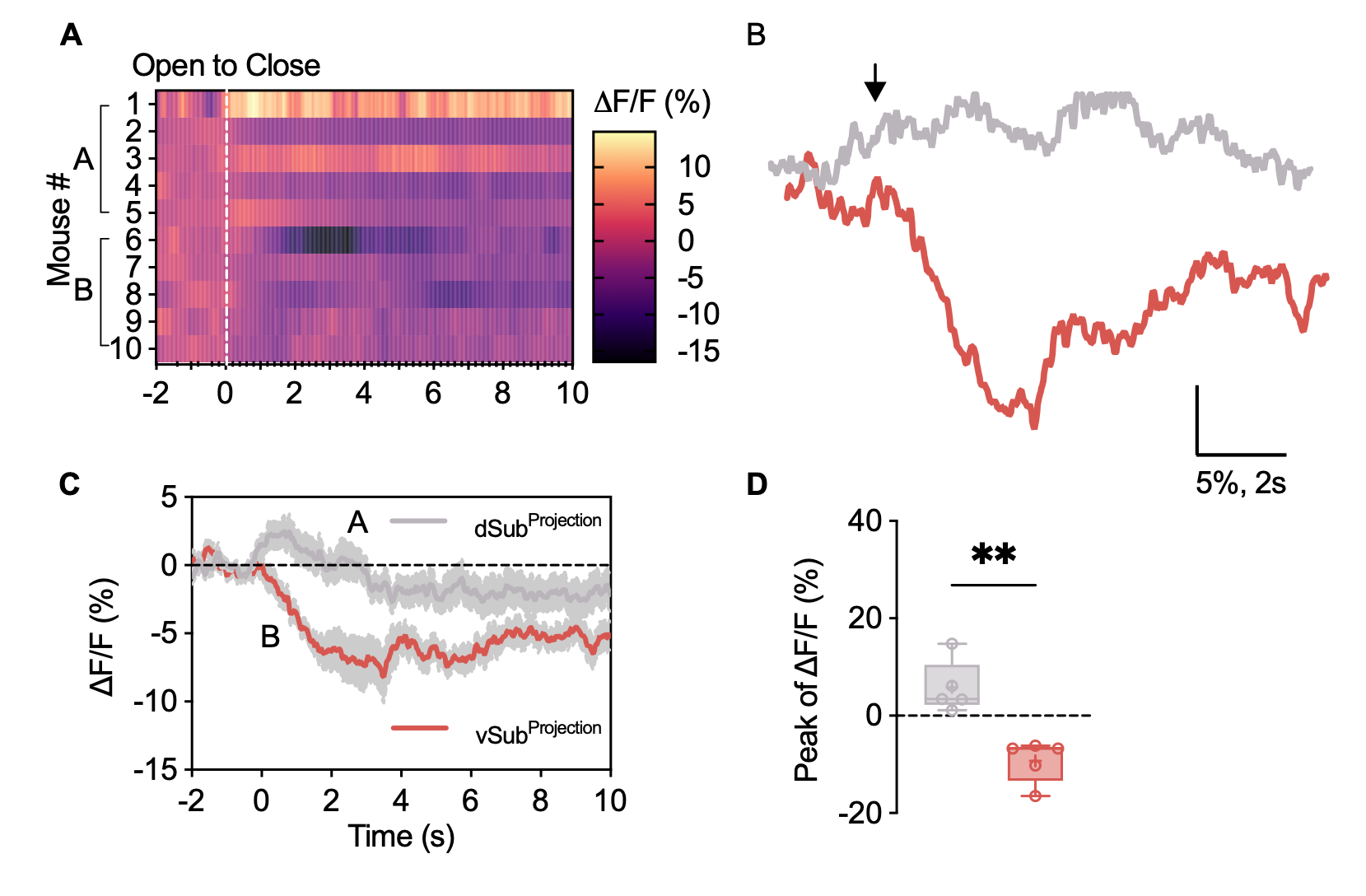

**Supplemental Figure 5.** Calcium fiber photometry during EPM test

**(A)** Heatmap of the Ca^2+^ fluorescence intensity during the transition from the open arms to the closed arms. **(B)** Representative Ca^2+^ activity during the transition from the open to the closed arms. **(C)** Average ∆F/F of Ca^2+^ recorded in the dSub and vSub. **(D)** Statistical analysis of the peak Ca^2+^ activity. n = 5 per group; unpaired t test

| **Table 1.** Statistical details | | | | | | |
| --- | --- | --- | --- | --- | --- | --- |
| **Figure** | **Panel** | **Statistical method** | **t/U/F value** | **df/sum of ranks** | **95% CI** | **p value** |
| **1** | **C** | Unpaired t test | 3.811 | 21 | 100.0 to 340.2 | 0.001 |
|  | **F** | Two-way ANOVA | Interaction, Time, Naïve vs. CSDS =15.27, 7.776, 15.27 | 1,72 | - | <0.001, 0.007, <0.001 |
| **2** | **D** | Unpaired t test | 5.15 | 6 | 28.1 to 78.9 | 0.002 |
|  | **E** | Unpaired t test | 0.614 | 18 | -7.227 to 13.19 | 0.547 |
|  | **F** | Mann Whitney test | 33 | 88,122 | - | 0.218 |
|  | **G** | Unpaired t test | 3.386 | 14 | -38.78 to -8.703 | 0.004 |
|  | **H** | Mann Whitney test | 7.5 | 108.5,62.5 | - | 0.002 |
| **3** | **C** | One-way ANOVA | 39.9 | 4,10 | - | <0.001 |
|  | **F** | One-way ANOVA | 151 | 2,6 | - | <0.001 |
|  | **G** | One-way ANOVA | 16.5 | 2,6 | - | 0.004 |
|  | **H** | One-way ANOVA | 14.08 | 2,6 | - | 0.005 |
| **4** | **B** | Two-way ANOVA | Interaction, Time, Control vs. Foot-shock =1.903,5.155,4.886 | 1,38 | - | 0.176,0.029,0.033 |
|  | **D** | Unpaired t test | 2.59 | 7 | -306 to -25.0 | 0.025 |
|  | **F** | Unpaired t test | 8.793 | 7 | 51.22 to 88.90 | <0.001 |
|  | **I** | Unpaired t test | 4.24 | 27 | -13.7 to -4.77 | <0.001 |
|  | **J** | Unpaired t test | 4.31 | 27 | -67.5 to -24.0 | <0.001 |
|  | **K** | Mann Whitney test | 58 | 272,163 | - | 0.041 |
|  | **L** | Unpaired t test | 3.532 | 26 | -0.1930 to -0.05100 | 0.002 |
| **5** | **L** | Unpaired t test | 2.696 | 8 | 1.372 to 17.58 | 0.027 |
|  | **M** | Unpaired t test | 2.631 | 8 | 2.332 to 35.45 | 0.03 |
|  | **Q** | Unpaired t test | 1 | 9 | -4.897 to 12.66 | 0.343 |
|  | **R** | Unpaired t test | 0.422 | 9 | -51.03 to 74.45 | 0.683 |
| **6** | **C** | Two-way ANOVA | Interaction, Time, Column =0.266,0.109,5.79 | 30,319 |  | 1,1,<0.001 |
|  | **E** | Two-way ANOVA | Interaction, Naïve vs. CSDS, mCherry vs. Gi =0.135,0.005,0.615 | 1,29 |  | 0.716,0.944,0.439 |
|  | **F** | Two-way ANOVA | Interaction, Naïve vs. CSDS, mCherry vs. Gi =11.5,2.998,3.315 |  |  | 0.002,0.094,0.079 |
|  | **G** | Two-way ANOVA | Interaction, Naïve vs. CSDS, mCherry vs. Gi =18.5,11.4,2 |  |  | <0.001,0.002,0.168 |
| **S2** | **B** | Unpaired t test | 0.62 | 18 | -1316 to 2427 | 0.54 |
|  | **C** | Unpaired t test | 0.639 | 14 | -746.6 to 1380 | 0.533 |
|  | **D** | Unpaired t test | 3.89 | 27 | -2315 to -715 | <0.001 |
|  | **E** | Unpaired t test | 3.079 | 27 | -1173 to -234.8 | 0.005 |
| **S6** | **D** | Unpaired t test | 4.893 | 8 | -22.15 to -7.961 | 0.001 |
